# Automated Annotation and Validation of Human Respiratory Virus Sequences using VADR

**DOI:** 10.1101/2025.08.07.669219

**Authors:** Jeffrey Furlong, Stephanie Goya, Eric P. Nawrocki, Vincent Calhoun, Eneida Hatcher, Linda Yankie, Alexander L Greninger

## Abstract

Accurate annotation of viral genomes is essential for reliable downstream analysis and public data sharing. While NCBI’s Viral Annotation DefineR (VADR) pipeline provides standardized annotation and quality control, it only supports six viral groups to date. Here, we developed and validated 12 new reference sequence-based VADR models targeting key human respiratory viruses: measles virus, mumps virus, rubella virus, human metapneumovirus, human parainfluenza virus types 1–4, and seasonal coronaviruses (229E, NL63, OC43, HKU1). Model construction was guided by a comprehensive analysis of intra-species genomic and phylogenetic diversity, enabling the development of genotype-specific models associated with reference genomes that defined expected genome structure and annotation. Models were trained on 5,327 publicly available complete viral genomes and tested on 372 viral genomes not yet submitted to GenBank. VADR passed 96.3% of publicly available viral genomes and 98.1% of viral genomes not in the training set, correctly identifying overlapping ORFs, mature peptides, and transcriptional slippage as well as genome misassemblies. VADR detected novel viral biology including the first reported HCoV-OC43 NS2 knockout in a human infection and novel G and SH coding sequence lengths in human metapneumovirus. These VADR models are publicly available and are used by NCBI curators as part of the GenBank submission pipeline, supporting high-quality, scalable viral genome annotation for research and public health.

## Introduction

Viral genomic sequencing plays a critical role in detecting emerging viral variants, informing diagnostic development, and shaping epidemiological surveillance and public health response. Decades of influenza virus sequencing have enabled effective surveillance, evolutionary forecasting for updating vaccines, and antiviral resistance tracking (1). The COVID-19 pandemic resulted in dramatic increases in viral genomic sequencing capacity for tracking SARS-CoV-2 variant evolution, which has since catalyzed increased sequencing of other respiratory viral pathogens (Figure 1A). Spurred by the rollout of new vaccines and prophylactic monoclonal antibodies, respiratory syncytial virus (RSV) genome submissions to International Nucleotide Sequence Database Collaboration (INSDC) databases have surged 325% in 2024 compared to 2023, reaching 6,402 entries. Genomics-driven viral surveillance is now an expectation of public health and regulatory authorities and will continue to expand as more medical interventions for viral infections are generated (2).

**Figure 1.**
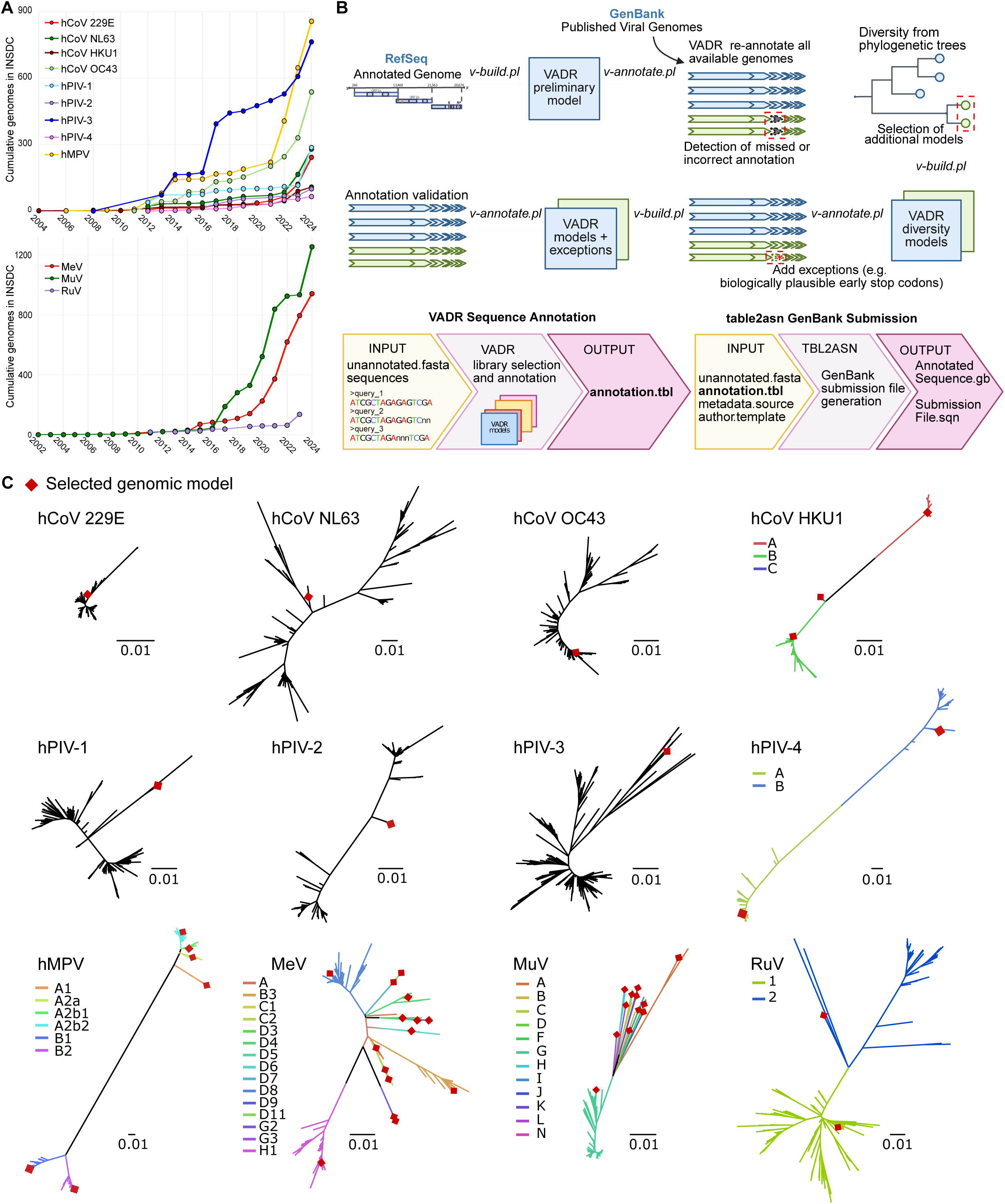
VADR model development of human respiratory viruses. A) Annual increase in complete genome submissions to NCBI GenBank for human respiratory viruses. The upper plot shows submissions for seasonal respiratory viruses: human metapneumovirus (hMPV), human coronaviruses (HCoV-229E, -HKU1, -NL63, - OC43), and human parainfluenza viruses (types 1–4). The lower plot displays submissions for vaccine-preventable respiratory viruses: measles virus (MeV), mumps virus (MuV), and rubella virus (RuV). B) Workflow for VADR model construction. RefSeq prototype genomes were used for initial model generation, except for HCoV-NL63 due to incomplete mature peptide annotations (see Methods). Following initial annotation, iterative refinement steps included phylogenetically informed model selection and the incorporation of alternative genomic features to optimize annotation performance and achieve >95% success across datasets. C) Maximum-likelihood phylogenetic trees of complete genomes analyzed for each virus. Red diamonds indicate genome sequences selected for inclusion in the final VADR model libraries. In cases where multiple models were selected, tree branches are color-coded to represent distinct phylogenetic clades/genotypes.

As the volume and breadth of viral sequencing increases, new challenges have emerged related to sequence quality control, standardization, and accurate genome annotation (3). While bioinformatic tools for consensus viral genome generation are now widely available – both as local software or web-based platform – challenges persist in verifying whether the generated genomes are, indeed, accurate (3,4). Genome annotation is a critical part of sequence quality control and involves identifying and labeling functional elements such as protein coding sequence (CDS), mature peptides, and regulatory motifs. This process defines the coordinates of each feature and assigns biological meaning, transforming raw sequence data into interpretable, structured information essential for tracing amino acid changes in emerging viral variants and linkage with databases such as UniProt or National Center for Biotechnology Information (NCBI) Protein or Gene. Despite its importance, genome annotation is often time-consuming and challenging to scale across viral species, highlighting the need for robust automated pipelines that ensure both efficiency and quality control (5,6). While a number of automated viral annotation pipelines have been generated to date, they generally lack the sequence quality control steps that allow them to integrate seamlessly with INSDC or NCBI databases (5–9).

To address these challenges, NCBI developed Viral Annotation DefineR (VADR), a software pipeline which automates the validation and annotation of viral genome submissions to NCBI GenBank public database (10). VADR employs curated genomic models to classify sequences by comparing them to a reference library of prototype homology models and transfers the annotation to the query sequence based on an alignment to the best-matching model. Predicted proteins are confirmed using BLASTX. During the annotation process, nucleotide sequences undergo 70 different quality control checks, including frameshifts, premature stop codons, reverse complement sequences, duplicate regions, low similarity scores, missing start codons, and indefinite annotations. As such, VADR can automate portions of the sequence quality control performed by GenBank curators and also serve as an ideal final step in user-designed consensus genome assembly pipelines to detect misassemblies and identify novel viral features. Because of the complexity of viral biology, VADR models must be validated for each viral species and have so far been limited to only six species, including norovirus, dengue virus, SARS-CoV-2, influenza virus, RSV, and monkeypox virus (11,12).

Here, we present an expanded set of VADR models for seasonal and vaccine-preventable respiratory viruses, including seasonal human coronaviruses (HCoV-229E, - NL63, -OC43, and -HKU1), human parainfluenza viruses (HPIV) types 1-4, human metapneumovirus (hMPV), measles virus (MeV), mumps virus (MuV), and rubella virus (RuV). We selected representative genomes using a phylogenetic framework and a thorough analysis of genotype-based genomic diversity, genome completeness, and rigor of prior annotation for each species. These models were tested against both publicly available GenBank genomes and viral genomes generated by the University of Washington Virology Laboratory not in the training set as part of other projects (13,14). The expanded VADR annotation models are freely available via GitHub (https://github.com/greninger-lab/vadr-vscan-aglab) and are used by NCBI curators as part of the GenBank submission pipeline, supporting broader and standardized genome annotation of respiratory virus genomes.

## Materials and Methods

### Genomic Datasets from INSDC Databases

Complete viral genomes of seasonal human respiratory viruses and vaccine-preventable viruses without available VADR models – including human coronaviruses HCoV-229E, HCoV-NL63, HCoV-OC43, HCoV-HKU1, HPIV types 1-4, hMPV, MeV, MuV, and RuV – were downloaded from INSDC databases using the NCBI GenBank portal. Search criteria terms, genome length, download dates are provided in Supplementary Material. Sequences containing more than 10% ambiguous bases (“N”) were excluded from downstream analysis.

### Initial VADR Model Construction

VADR models were built iteratively: an initial model based on a single representative genome was used to annotate all genomes for each species, after which genomic and phylogenomic analyses guided the selection of additional representative genomes to capture known viral diversity. Initially, VADR models were mainly generated using prototype genomes from the NCBI RefSeq database (10). For HCoV-NL63, the RefSeq genome (NC_005831.2) lacked mature peptide annotations, so NCBI GenBank accession ON553997.1 was used instead. In addition, RuV has two representative RefSeq genomes (NC_001545.2 and NC_076948.2) which include the complete polyprotein but lack a poly(A) tail at the 3’ untranslated region (UTR). Thus, to establish a RuV model with a full genome, NCBI GenBank accession AB928205.1 was used for the preliminary model. For each species, the initial model was created using the VADR v-build.pl script with default parameters using the --gb option to parse the prototype genomes from their NCBI accession number (Figure 1B).

The initial models were used to annotate all available complete genomes in NCBI GenBank using the VADR v-annotate.pl script, as described in the section “Annotation and Validation of Respiratory Virus Genomes using VADR”. This initial annotation round helped standardize genetic features and nomenclature across all genomes within each dataset. It also revealed recurrent VADR error alerts of biologically plausible genomes, likely due to insufficient diversity representation from a single RefSeq prototype genome.

### Final VADR Model Library Construction

To capture intra-species genomic diversity, iterative model testing was conducted by evaluating VADR annotation error alerts with updated model sets (Figure 1B). Phylogenetic analysis was also performed to ensure the models reflected phylogenetic diversity. Genome alignments were generated with MAFFT v7.221, and maximum-likelihood phylogenies were inferred using IQ-TREE v2 (15,16). Alignments and phylogeny visualizations were performed using Geneious Prime 2023.0.1 (https://www.geneious.com), AliView v1.28 (17) and Figtree v1.4.4 (http://tree.bio.ed.ac.uk/software/figtree/). Phylogenetic trees are available at https://github.com/greninger-lab/vadr-respiratory-virus.

Based on observed genomic and phylogenetic diversity (Figure 1C), representative genomes from distinct genotypes were selected and incorporated into the final model libraries. The longest available genomes were preferentially selected for the final prototypes to maximize feature coverage and genome completeness. Final VADR libraries were constructed using locally annotated genomes (-ingb option) to ensure standardized annotation with harmonized protein naming across the prototype sequence selected (Table 1). Final model libraries were used to re-annotate the datasets using VADR v-annotate.pl script. Annotation error alerts related to missing or truncated genes or protein annotation discrepancies were reviewed for recurrence. When errors were supported by either being detected in more than one genome or validation through reassembly from raw sequencing data available in NCBI SRA, alternate features were added to better inform the models of expected genomic diversity. Alternate features are strain-specific genetic elements that differ from the reference but must follow biological rules, for instance, a frameshift caused by a deletion that is corrected by a downstream mutation within the coding sequence.

**Table 1.** Summary of VADR model libraries and genome annotation performance. For each viral species, a detailed description of the annotation model library is provided. This includes the number of complete genomes analyzed, the NCBI GenBank accession numbers for the initial and final models, and the percentage of genomes that successfully passed VADR validation and annotation. In addition, biologically plausible alternative features (stop codons) identified by gene and number of occurrences are reported, as these were incorporated to improve the final library annotation success.

| Library* | Number of Genomes | Initial Model | % Initial Model Success | Final Models in the Library (Genotype) | Gene with Alternative CDS (Additional Features) | % Final Model Success |
| --- | --- | --- | --- | --- | --- | --- |
| hMPV | 821 | NC_039199.1 | 1.2 | NC_039199.1 (A1),<br>KF686742.1 (A2a),<br>KC562225.1 (A2b1),<br>KY474539.1 (A2b2),<br>OL794407.1<br>(A2b2_180dup),<br>OL794418.1<br>(A2b2_111dup),<br>MZ221202.1 (B1),<br>OL794387.1 (B2) | SH (6), G<br>(12) | 94.1 |
| HPIV-1 | 241 | NC_003461.1 | 94.1 | NC_003461.1 | P (1), L (1) | 97.5 |
| HPIV-2 | 113 | NC_003443.1 | 0.8 | NC_003443.1 | P (1), L (1) | 96.4 |
| HPIV-3 | 847 | NC_001796.2 | 0.2 | NC_001796.2 | P (1), HN<br>(1) | 96.2 |
| HPIV-4 | 70 | NC_021928.1 | 2.8 | LC706550.1 (A),<br>EU627591.1 (B) | F (1), HN<br>(2) | 98.5 |
| HCoV<br>229E | 219 | NC_002645.1 | 12.3 | KY996417.1 | None | 96.3 |
| HCoV<br>NL63 | 259 | ON553970.1 | 33.2 | PQ187621.1 | M (3) | 98.4 |
| HCoV<br>OC43 | 382 | NC_006213.1 | 1.5 | MT118678.1 | E (2), I (1) | 96.8 |
| HCoV<br>HKU1 | 123 | NC_006577.2 | 35.7 | NC_006577.2 (A),<br>AY884001.1 (B),<br>DQ415897.1 (C) | None | 77.2 |
| MuV | 1141 | NC_002200.1 | 8.6 | MH426702.1 (A),<br>MK279727.1 (B),<br>KX953297.1 (C),<br>MW819866.1 (D),<br>DQ649478.1 (F),<br>OR822028.1 (G),<br>KY969483.1 (H),<br>AY309060.1 (I),<br>KF878079.1 (J),<br>MT238690.1 (K),<br>KF878081.1 (L),<br>AY685921.1 (N) | N (1), SH<br>(2) | 99.8 |
| MeV | 940 | NC_001498.1 | 93.7 | AF266288.2 (A),<br>OR290098.1 (B3),<br>LC655230.1 (C1),<br>MG912589.1 (C2),<br>MN017369.1 (D11),<br>AB481088.1 (D3),<br>OP236009.1 (D4),<br>JN635406.1 (D5),<br>DQ227319.1 (D6),<br>JN635410.1 (D7),<br>PP101943.1 (D8),<br>KY969476.1 (D9),<br>MG912591.1 (G2),<br>KC164758.1 (G3),<br>MZ442631.1 (H1) | H (1), V (1),<br>F (2) | 94.4 |
| RuV | 171 | NC_001545.2 | 99.8 | MK780811.1 (clade 1),<br>DQ388279.1 (clade 2) | None | 100 |

(\*) hMPV, human metapneumovirus, HPIV-1, human parainfluenza virus 1, HPIV-2, human
parainfluenza virus 2, HPIV-3, human parainfluenza virus 3, HCoV, human coronavirus,
MuV, mumps virus, MeV, measles virus, RuV, rubella virus.

A complete list of final prototype genomes used per species is available in Table 1. The GitHub websites of the 12 new libraries of VADR models are listed in Supplementary Material. We provide a Docker containerized application to automate the selection of the appropriate model library, sequence annotation and validation as well as generation of NCBI GenBank submission files, available at https://github.com/greninger-lab/vadr-vscan-aglab (Figure 1B).

### Annotation and Validation of Respiratory Virus Genomes using VADR

Genome annotation was performed in batch using the VADR v-annotate.pl script with the virus-specific built model libraries and a FASTA sequence file as input. For all viruses, the -r option was used to temporarily replace ambiguous stretches of Ns with reference-matching nucleotides to facilitate feature identification. Additional virus-specific parameters were applied as needed:

For measles virus (MeV) and mumps virus (MuV), the –indefclass threshold was adjusted to 0.01 and 0.025, respectively, to improve discrimination between closely related prototype genomes to facilitate genotyping. This parameter defines the minimum bits-per-nucleotide difference between the top two models to trigger an “indefinite classification” alert and defaults to 0.03. Due to high similarity of the 15 MeV and 12 MuV genotype reference genomes (>95% average pairwise percent identity), indefinite classification alerts were more frequent compared to other VADR virus model libraries. Thresholds values were optimized to balance sensitivity to potential misclassifications with reduction of spurious alerts when classifications matched.

For human coronaviruses, the --glsearch option was used, consistent with the available SARS-CoV-2 VADR model. This BLASTN-seeded strategy uses local alignment to determine seed region, with only unaligned regions processed using the memory-efficient glsearch program. Compared to the default cmalign method, this approach reduced peak memory usage from 64Gb to 2Gb and substantially decreased run times (12). In addition, the --alt_pass discontn option was used for HCoV-HKU1 and HCoV-OC43 models to allow annotation despite discontinuous similarity alerts caused by sequence features such as short repeats in the HCoV-HKU1 ORF1ab nsp3 peptide region or alternative transcription regulatory leader-like sequences in HCoV-OC43 (18,19).

A genome was considered to pass VADR annotation if all expected annotation features were correctly identified and no other fatal alerts were reported. Genome validation rate was calculated as: (number of VADR fully annotated genomes / total genomes analyzed) × 100%.

### Validation of Final VADR Models Using New Viral Genomes

Remnant clinical samples positive for seasonal human respiratory viruses from the University of Washington Medicine Virology Laboratory were sequenced as previously described, or using QIAseq xHYB Respiratory Panel hybridization capture, following manufacturer’s recommended protocol (20). DNA libraries were sequenced as 1x100bp or 2x150bp reads on Illumina NovaSeq6000 or NextSeq2000 platforms. The use of deidentified specimens was approved by the University of Washington Institutional Review Board with a consent waiver (protocol no. STUDY00000408).

Consensus genome assemblies were generated using REVICA-STRM pipeline (https://github.com/greninger-lab/revica-strm), including reads quality filtering and mapping to reference genomes with a minimum variant call depth of 5X. Genomes were annotated with the VADR v-annotation.pl script as described in the section above. NCBI GenBank accession numbers are detailed in Supplementary Table 1.

## Result

### VADR Model Generation and Annotation Performance

The VADR model library for each virus species was developed and iteratively refined by evaluating annotation error alerts and incorporating representative genomes, alternate features, and exceptions annotations. While model construction for a reference sequence and its features is automated with the v-build.pl script, addition of alternative features and exceptions requires adding new proteins to the BLASTX library files and manual editing of the model info file as described in the VADR documentation. The objective was to maximize genome validation rates while ensuring accurate feature annotation and compatibility with biologically plausible genetic variation.

### Seasonal Respiratory Viruses

#### Human seasonal coronaviruses (HCoV)

Initial annotation success using single RefSeq-based models was low for all four seasonal human coronaviruses, with validation rates of 12.3% for HCoV-229E, 33.2% for HCoV-NL63, 35.7% for HCoV-HKU1, and 1.5% for HCoV-OC43. These low rates underscored the high genomic diversity of circulating HCoV and the limitations of using a single prototype genome for annotation. Refinement of models by selecting more representative genomes and incorporating sequence-specific features markedly improved performance.

For HCoV-229E, using a clinical isolate (reference genome GenBank accession number KY996417.1) instead of the lab-adapted RefSeq genome (NC_002645.1) raised the success rate from 12.3% to 96.3%. The lab-adapted RefSeq genome carries a two-nucleotide deletion that splits ORF4 into ORF4a and ORF4b, whereas clinical strains typically encode a single ORF4, likely reflecting adaptation to in vitro growth conditions (21). Similarly, for HCoV-NL63, substitution of a divergent genome (ON553970.1) with a representative genome (PQ187621.1) that is phylogenetically closer to circulating strains and includes more complete 5’ and 3’ UTRs improved the annotation success to 98.4% of genomes (Figure 1C). In this updated model, three alternative Membrane (M) gene variants were added to account for insertions of 6, 9, or 21 nucleotides observed in 44 genomes (PP591771.1, PQ187626.1, ON553970.1, respectively).

For HCoV-OC43, annotation performance improved when the laboratory strain genome ATCC VR-759 (NC_006213.1) was replaced with a clinically derived genome (MT118678.1), which showed higher similarity across the Envelope (E) and Spike (S) proteins (22). Additional model features were incorporated to reflect variation in the stop codon for the E gene, either extending the protein by one amino acid due to a frameshift (single nucleotide deletion in position 28369 observed in 48 genomes, KF530064.1) or four amino acids (28369-28374del reported in 72 genomes, KF530068.1). Similarly, a truncation of 147 amino acids in the Internal (I) protein was added as alternate feature due to an early stop reported in 7 genomes with the substitution c.29332C>U (NC_006213.1).

HCoV-HKU1 model performance improved with the addition of genotype-specific models representing genotypes A (NC_006577.2), B (AY884001.1), and the recombinant genotype C (DQ415897.1). A feature exception was included in this library to account for the variable tandem copies of a 30-nt repeat at the N-terminus of nsp3 inside the ORF1a gene, raising the annotation success from 35.7% to 77.2%. (19). During the annotation process, we identified 27 HCoV-HKU1 genomes that failed VADR validation due to low similarity compared to the model in the Hemagglutinin-esterase (HE) and Spike genes, largely attributed to a high proportion of Ns, reaching up to 38% in some cases. We hypothesized that these gaps resulted from assemblies based on reference genome from NCBI RefSeq (which is genotype A) that did not fully capture genotypes B and C specific genetic signatures. Following communication with the original submitters and reanalysis of the raw sequencing reads, 17 assemblies were successfully updated, passed VADR validation, and were resubmitted to GenBank. The remaining 10 genomes could not be reassembled and VADR currently annotates the affected genes as miscellaneous features.

#### Human parainfluenza viruses (HPIV)

Initial model performance varied across HPIV species not related with overall sequence similarity issues but due to the presence of a significant number of alternate features. The HPIV-1 model achieved 94.1% success with the single RefSeq model, which improved to 97.5% after accounting for mutations in the P and L genes that extended or truncated the proteins. Specifically, two genomes exhibited a mutation in the P stop codon (c.3548U>C, PP334213.1) extending the protein by two amino acids, and two genomes exhibited a mutation in the L gene (c.15438C>U, PQ117904.1) causing an early stop codon truncating the protein by one amino acid.

Initial models for HPIV-2 and HPIV-3 showed low validation rates of 0.8% and 0.2%, respectively. For HPIV-2, the annotation success rate was improved by recognizing a compensatory frameshift event in the P gene, involving an insertion at position 2938 and a nearby deletion at position 2945, present in 112 genomes (OQ848525.1). Another compensatory frameshift was modeled in the L gene, including a nucleotide deletion in position 13987 and a two-nucleotide deletion downstream (14017-14018del) exhibited in 107 genomes (PP334183.1). For HPIV-3, early stop codon mutations at the end of P and HN genes were added to the model as alternate features. Specifically, the mutation c.3590C>U in the P gene present in 735 genomes (AB736166.1) truncated the protein by one amino acid, and the mutation c.8522C>U in the HN gene detected in 830 genomes (AB736166.1) caused a two-amino acid truncation.

For HPIV-4, the RefSeq-based model HPIV-4A M-25 (NC_021928.1) was replaced with a clinically derived genome (LC706550.1), and a representative model for HPIV-4B was added (EU627591.1). Two alternate features for HPIV-4A were detected in three genomes (AB543336.1): a premature stop codon (c.6863C>U) in the F gene truncating the protein by one amino acid and a two-nucleotide insertion (9284–9285) in the HN gene causing frameshift that extended the protein by 5 amino acids. For HPIV-4B, a compensated-frameshift event in the HN gene was observed in two genomes and it was incorporated (single nucleotide deletion in position 8371 followed by an insertion in position 8414, JQ241176.1). These updates allowed the refined model to successfully annotate 98.5% of the HPIV-4 genomes.

#### Human metapneumovirus (hMPV)

The initial hMPV annotation model based solely on the RefSeq prototype genome (NC_039199.1) successfully annotated only 1.2% of genomes. Most annotation error alerts were associated with low similarity in SH and G genes, underscoring the high genetic variability of these regions. To address this, we expanded the model incorporating reference genomes from all six known hMPV lineages: A1 (NC_039199.1), A2a (KF686742.1), A2b1 (KC562225.1), A2b2 (KY474539.1), B1 (MZ221202.1), and B2 (OL794387.1) (23). In addition, two variants from the A2b2 lineage, each containing nucleotide duplication within the G gene, were included: one with a 111-nt duplication (OL794418.1) and another with a 180-nt duplication (OL794407.1) (24,25). The incorporation of alternative gene features further accounted for lineage-specific diversity, ultimately improving genome annotation coverage to 93.5% (Figure 1C).

Lineage-specific variations affecting SH and G gene annotations were particularly common, requiring the addition of alternative features. In the A1 lineage, a substitution in SH gene stop codon extended the protein by six amino acids observed in five analyzed genomes (KC403977.1). In addition, a 20-nt deletion at position 6898 in the G gene caused a frameshift that introduced early stop codons at positions 6924 (KC562241.1) and 6951 (PP086007.1), truncating the G protein by 17 and 8 amino acids, respectively. Within lineage A2a, multiple mutations affected both the SH and G genes. At position 5976, stop codon mutations produced two SH proteins variants: one extended by two amino acids observed in two genomes (PP086006.1), and another by four amino acids observed in four genomes (AB503857.1). Another mutation at position 6841(c.6841U>C) in the G gene stop codon led to extended proteins of varying lengths: two amino acids (observed in 18 genomes, OL794370.1), seven amino acids (15 genomes, KJ627413.1), eleven amino acids (10 genomes, AB503857.1), and twelve amino acids (2 genomes, KJ627398.1). In the A2b1 lineage, a c.6857C>U mutation introduced a premature stop codon in the G gene, resulting in a truncated protein lacking the final two amino acids (47 genomes, OL794373.1). Conversely, the c.6867U>C mutation caused a nine-amino acid extension of the G protein (14 genomes, OL794361.1).

The A2b2 lineage included variants of the 111-nt duplicated G gene, which also exhibited truncation caused by premature stop codons. Specifically, the c.6973C>U mutation truncated the G protein by nine amino acids (4 genomes, LC769215.1), while c.6994C>U led to a two-amino acid truncation (7 genomes, MN745084.1).

In the B1 lineage, two distinct mutations affected the SH gene stop codon. A c.5947U>A mutation extended the SH protein by two amino acids (8 genomes, OY757729.1), and a 3-nt deletion overlapping the SH stop codon (positions 5948-5950) resulted in an 18-amino acid extension (63 genomes, OP904074.1). In addition, two G gene mutations caused protein truncations: c.6852C>U shortened the protein by ten amino acids (58 genomes, OP904074.1), and c.6858C>U by eight amino acids (6 genomes, OP904104.1). Finally, in B2 lineage, a c.5999U>C mutation in the SH gene stop codon extended the protein by 23 amino acids (3 genomes, KC562229.1).

#### Testing VADR Models with New Data for Seasonal Respiratory Viruses

To independently evaluate the refined models, we took advantage of ongoing sequencing projects related to these viruses in our laboratory. We ran our newly generated VADR models on 372 viral genomes sequenced from clinical respiratory virus samples at the University of Washington Virology Laboratory. These included genomes from hMPV (n= 230), HPIV-1 (n= 17), HPIV-2 (n= 25), HPIV-3 (n= 11), HPIV-4 (n= 32), HCoV-229E (n= 13), HCoV-NL63 (n= 8), HCoV-OC43 (n= 17), and HCoV-HKU1 (n= 19). None of these genomes were part of the training datasets used for VADR model construction described above.

Of the 372 viral genomes, 365 genomes (98.1%) were validated and fully annotated by VADR. Of the seven sequences that returned alerts, four had varying hMPV SH and G coding sequence length and were supported by sequencing reads, while two HCoV-HKU-1 sequences were corrected prior to submission and one HCoV-OC43 genome illustrated novel biology.

In one HCoV-OC43 genome (PV685489.1), VADR reported a premature stop codon at amino acid 21 of the NS2 accessory protein, which was confirmed upon preparing new libraries and resequencing the specimen. NS2 has been shown to be a non-essential gene in HCoV-OC43 in cell culture and mouse infections, but never in humans (26,27). One HCoV-HKU1 genome (PV685464.1) showed misalignments in a poly(U) homopolymer region leading to a spurious frameshift while another sequence had a non-sense mutation in ORF1a (PV685465.1), which were both resolved upon preparing new libraries and resequencing the specimens.

Among hMPV sequences, two genomes had stop codon shifts in the G gene, with supporting evidence from read coverage. Specifically, hMPV A2b2 genome PV052163.1 had a frameshift in the G gene stop codon supported by 247X coverage that led to 4 amino acids truncation and PV052138.1 had a two-amino acid truncation in G based on a c.6857C>U mutation supported by 10X depth. Two additional genomes exhibited ambiguous nucleotide regions (Ns) that resulted in partial annotations of M2 and SH coding regions in PV052203.1 and the G gene in PV022071.1.

### Vaccine-Preventable Viruses

#### Measles virus (MeV)

The RefSeq-based initial model for MeV (NC_001498.1) achieved 93.7% annotation success across 940 GenBank MeV genomes. To support genotyping, we expanded the model library to include representative genomes for all WHO-designated measles genotypes spanning clades A through H (Table 1, Figure 1C) (28). This increased the annotation success to 94.4% and enabled 100% genotype classification accuracy. Genotype-specific alternate features were added to account for alternate stop codons: a mutation in the stop codon causing four amino acid extension in the H protein observed in genomes within genotypes B3 (ON024067.1), C1 (LC655227.1), D3 (AB254456.1), and D6 (MF775733.1); a mutation in the stop codon of the V protein of genotype C1 extending the protein by one amino acid (LC655227.1), and two mutations in the F protein including a single nucleotide deletion in position 7024 of genotype D3 causing an 11 amino acid mismatch and 16 amino acid truncation (GQ376026.1), and the substitution c.7036C>U causing a 24 amino acid premature truncation of genotype D6 (DQ227318.1).

#### Mumps virus (MuV)

The initial model using RefSeq accession NC_002200.1 successfully annotated only 8.6% of 1,141 MuV genomes. After incorporating representative genomes for all 12 known genotypes (A–D, F–L, N), annotation success increased to 99%, with full genotype resolution (Table 1, Figure 1C) (29). Genotype-specific variation in the NP and SH genes was incorporated into MuV models, including a substitution in the NP gene stop codon of genotype A extending the protein by 21 amino acids (JQ946552.1), a 7 amino acid truncation due to a premature stop codon (substitution c.6420G>A) in the SH gene of genotype G detected in five genomes (OK440543.1), and a substitution in the SH stop codon in 11 genomes extending the protein by 19 amino acids (MT880121.1).

#### Rubella virus (RuV)

Rubella virus has two representative genomes in RefSeq, corresponding to genomes of phylogenetic Clade 1 (NC_001545.2) and Clade 2 (NC_076948.2) (30). While these genomes include complete polyprotein sequence and 3’UTR, they lack a polyA tail. To establish a model with confirmed complete genome, accession number AB928205.1 (Clade 2) was selected, resulting in the annotation of 170 out of 171 genomes (99.8%), though clade classification was absent. Final models incorporated representative genomes from both Clade 1 (MK780811.1) and Clade 2 (DQ388279.1). The Clade 2 genome was updated to select genotype 2C, more closely related to Clade 1, to ensure accurate classification and prevent genotype misidentification if selecting a model genetically more divergent. The final library models enabled 100% annotation success and accurate clade assignment.

## DISCUSSION

The rapid expansion of viral genome sequencing for surveillance, outbreak response, and antiviral and vaccine development has underscored the need for robust and standardized annotation tools. The VADR framework has been a cornerstone of the quality control pipeline for NCBI GenBank for select viruses such as influenza virus and SARS-CoV-2 (11,12). Here, we used phylogenetic and genomic diversity of 12 common human respiratory viruses to expand VADR libraries from 6 to 18 viral species, encompassing most seasonal and vaccine-preventable respiratory pathogens. In addition, we validated these models on 372 viral sequences generated by our lab prior to submission to NCBI GenBank. This work addresses a critical gap in scalable, automated annotation of viral genomes essential for public health, particularly as volumes of submitted viral sequences continue to grow exponentially (31).

Initial models for HCoVs (229E, NL63, OC43, HKU1), HPIV 1–4, hMPV, MeV, MuV, and RuV based on lab-adapted or outdated references from NCBI RefSeq failed to fully annotate 70% of all analyzed genomes, reflecting the reference bias pervasive in viral genomics (4). To mitigate these limitations, we employed phylogenetically representative genomes that capture intraspecies circulating diversity and incorporated alternate features to accommodate biologically plausible variants, especially in highly variable surface glycoproteins. These alternate features included lineage-specific stop codons, nucleotide insertions, deletions and duplications observed in more than one analyzed genome.

The improved VADR libraries achieved high annotation success rates (>95% for most of virus-specific libraries) during the reannotation of 5,327 NCBI GenBank records. In addition, 197 genomes (3.7%) triggered error alerts from which 109 were due to insertions or deletions causing frameshifts and unexpected CDS boundaries observed in individual genomes.

Emerging approaches based on artificial intelligence trained on homology patterns are promising to characterize newly discovered viruses (32,33). However, accuracy and consistency are mandatory in public health surveillance, and the VADR-curated, rules-based approach continues to provide a reliable methodology for NCBI GenBank submissions. Integration of VADR into the NCBI GenBank submission workflow currently automates the detection of >70 common issues, substantially reducing the burden on curators during pre-release screening (10). Moreover, local use of VADR as a pre-submission quality control tool can help reduce misassemblies before submission. For instance, during quality control and annotation of 372 viral genomes prior to submission to GenBank, we identified genome assembly issues – such as a poly(U) region insertion in HCoV-HKU1 causing a frameshift – that can distort evolutionary analyses, as previously reported for SARS-CoV-2, RSV, and hMPV (3,34).

Importantly, VADR also helped to detect biologically significant variants such as the premature stop in HCoV-OC43 NS2 coding sequence supported by sequencing reads. NS2 is an accessory gene that codes for a phosphodiesterase that antagonizes RNase L activation and helps HCoV-C43 evade the host innate immune response (35). Although NS2 has been shown to be non-essential for HCoV-OC43 replication in cell culture and in mice (26), this is the first report of an NS2-truncating mutation in a human HCoV-OC43 infection. While accessory genes in coronaviruses are generally evolutionary advantageous and maintained in the genome, there are numerous reports of accessory gene loss in other coronaviruses (36–38).

Despite these advances, several limitations remain. Annotation success with VADR is sensitive to genome completeness and ambiguity. While VADR can handle sequences as short as 100–150 nucleotides in conserved regions, performance declines in highly variable regions or in sequences with stretches of ambiguous bases (Ns) of unexpected length (10). Annotation is most impacted when Ns occur in conserved nucleotide signatures or when the length of the N-stretch is not a multiple of three, leading to failure to pass protein validation. In addition, underrepresented genotypes (e.g., MeV genotypes B2, D2, D10, and H2) lack sufficient sequence data for robust modeling, and emergent strains may harbor novel features not captured by existing reference sequences, highlighting the need for ongoing model updates. Future efforts should prioritize collaborative model refinement leveraging community feedback. In addition, as our clinical laboratory seldom detects or sequences MeV, MuV, and RuV specimens, we did not provide newly sequenced isolates to validate the VADR models generated here. Nonetheless, the MeV models generated here are currently being beta-tested by NCBI GenBank curators on newly submitted MeV sequences and have not had issues reported to date.

Many challenges in viral genome annotation remain the same as those identified over 15 years ago, including the complexity of diverse gene structures that defy universal prediction methods, the risk of propagating errors from inaccurately annotated reference genomes, and the lack of standardized and consistent protein nomenclature, which leads to confusion and inaccuracies within databases (39). The expansion of VADR library models to cover all key respiratory viruses represents a critical step toward standardizing viral genome annotation. Its dual role in submission validation and quality control addresses urgent needs in public health genomics, where rapid data sharing must not come at the cost of accuracy.

## AUTHOR CONTRIBUTIONS

Conceptualization: JF, EPN, ALG. Formal analysis: JF, SG. Methodology: JF, ALG. Visualization: JF, SG. Validation: EPN, VC, EH, LY. Writing – original draft: JF, SG, ALG. Writing—review & editing: all authors.

## ACKNOWLEDGEMENTS

We thank all contributors to the NCBI GenBank database for sharing viral nucleotide sequences and metadata.

