## Supplementary Material for "Automated Annotation and Validation of Human Respiratory Virus Sequences using VADR"

**MATERIAL AND METHODS**

**Viral Genome Datasets from NCBI GenBank.**

Complete genomes for each viral species were downloaded from NCBI GenBank. NCBI metadata was reviewed to confirm viral genomes were obtained from human samples. Complete genomes with more than 10% ambiguous nucleotides (“N”) were removed before further analysis.

For each viral species the search terms and filters used to download the data were:

| Virus Name | Sequence Length | Database Accession Date |
| --- | --- | --- |
| Human metapneumovirus | 13-14 kb | August 05, 2024 |
| Human parainfluenza virus 1 | 15-16 kb | November 04, 2024 |
| Human parainfluenza virus 2 | 15-16 kb | October 09, 2024 |
| Human parainfluenza virus 3 | 14.4-16 kb | October 07, 2024 |
| Human parainfluenza virus 4 | 16-17.5 kb | October 07, 2024 |
| Human coronavirus 229E | 25-27.5 kb | November 25, 2024 |
| Human coronavirus NL63 | 25.5-28 kb | December 05, 2024 |
| Human coronavirus OC43 | 28.5-32 kb | December 05, 2024 |
| Human coronavirus HKU1 | 29-32 kb | December 05, 2024 |
| Mumps orthorubulavirus | 15-16 kb | April 15, 2025 |
| Measles morbillivirus | 15-17 kb | March 25, 2025 |
| Rubella virus | 9.5-10 kb | May 15, 2025 |

**GitHub of New Developed VADR Models.**

hMPV v1.01: https://github.com/greninger-lab/vadr-models-hmpv

HPIV (types 1–4) v1.0: https://github.com/greninger-lab/vadr-models-hpiv

HCoVs (229E, NL63, OC43, HKU1) v1.01: https://github.com/greninger-lab/vadr-models-hcov

MeV v1.01: https://github.com/greninger-lab/vadr-models-mev

MuV v1.0: https://github.com/greninger-lab/vadr-models-muv

RuV v1.0: https://github.com/greninger-lab/vadr-models-ruv

**Phylogenetic Trees of Datasets Downloaded from NCBI GenBank.**

Phylogenetic trees for each viral species are available at https://github.com/greninger-lab/vadr-respiratory-virus. Each tree tip contains the NCBI GenBank accession number. For viral species whose models included references of different genotypes, the genotype is detailed after the accession number.
